# Quantum machine learning for detection of sleep deprivation from EEG signals

**DOI:** 10.64898/2026.06.14.732153

**Authors:** Priyanshu Sarma-Sarkar, Rajkumar Saini, Partha Pratim Roy

## Abstract

Approximately 50% of the population in India is estimated to experience sleep-related disorders. Sleep deprivation is a prevalent condition that adversely impacts cognitive performance, neural functioning, and overall health. Electroencephalography (EEG) offers an objective means of capturing neural alterations associated with sleep loss, making it well-suited for automated detection frameworks. In this study, we explore the application of a Quantum Support Vector Machine and Hybrid Quantum Neural Networks to classify sleep-deprived and well-rested states using resting-state EEG signals.

A comprehensive feature extraction pipeline is employed, incorporating spectral band power, band ratios, Hjorth parameters, and functional connectivity measures. These features are subsequently encoded into quantum states to construct a quantum kernel, which is then utilized for classification. Model performance is evaluated under both epoch-level and subject-level data partitioning schemes.

The Hybrid Quantum Neural Network (HQNN) achieves the highest performance across both evaluation settings, attaining an accuracy of 96.88% at the epoch level and 81.25% at the subject level. The Quantum Support Vector Machine (QSVM) model achieves accuracies of 93.75% and 75.00% for epoch-level and subject-level evaluations, respectively. At subject-level and epoch-level evaluation, HQNN outperforms previously reported results (95.72% and 68.23%). Overall, these findings highlight the potential of quantum machine learning as a competitive approach for EEG-based sleep deprivation detection, with promising implications for real-world biomedical applications.

## I. Introduction

Sleep deprivation is a significant public health concern affecting cognitive performance, safety, and long-term physiological health. Acute sleep loss induces localized sleep, where specific brain regions enter sleep-like states despite wakefulness, leading to cognitive and motor lapses [1], [2]. These impairments are critical in high-stakes domains such as healthcare and transportation [3], and are associated with disrupted brain connectivity [4], reduced productivity [5], and increased impulsivity [6]. However, existing detection methods rely on subjective reports or laboratory-based assessments, limiting real-time applicability [7].

Electroencephalography (EEG) provides an effective modality for capturing neural signatures of sleep deprivation [8]. Sleep loss is consistently associated with increased low-frequency activity (delta, theta) and decreased high-frequency activity (alpha, beta) [2], [8], [9]. Region-specific effects include elevated slow-wave activity and reduced alpha connectivity in frontal regions [10], [11], and altered oscillatory patterns in parietal and occipital areas [8]. These findings highlight EEG’s suitability for objective detection of sleep deprivation.

Machine learning methods have been widely applied to EEG classification tasks. Ensemble models such as Random Forest, XGBoost, and LightGBM demonstrate strong performance [12–16], while SVMs remain effective for high-dimensional data [17]. Deep learning models (CNNs, LSTMs, Transformers) can learn spatiotemporal features but may over-fit on limited datasets [18–20]. Kumar et al. [21] showed that ensemble models generalize best for sleep deprivation classification, with key features including frontal theta activity and beta coherence.

Classical methods encode EEG features as classical vectors, potentially limiting the representation of complex neural structures. Quantum machine learning (QML) offers an alternative paradigm leveraging superposition and entanglement [22]. Quantum Machine Learning uses quantum feature maps to enable non-linear classification in high-dimensional spaces [23], and has shown promise in biomedical tasks [24–27].

Kumar et al. [21] have worked on the same problem of detection of sleep deprivation and the same dataset using classical machine learning and deep learning techniques. As shown in table I, without subject-level separation, CNN achieved the highest accuracy (95.72%), followed closely by XGBoost (95.42%), LightGBM (94.83%), and RF (94.53%). Under the more rigorous subject-level separation II, performance declined substantially across all models, with RF achieving the highest accuracy (68.23%), underscoring the difficulty of cross-subject generalization.

**TABLE I.** Classification performance of models at epoch-level separation from Kumar et al. [21].

| Model | Accuracy | F1-score |
| --- | --- | --- |
| LightGBM | $0.9483 \pm 0.0101$ | $0.9476 \pm 0.0095$ |
| Random Forest | $0.9453 \pm 0.0092$ | $0.9441 \pm 0.0096$ |
| XGBoost | $0.9542 \pm 0.0151$ | $0.9536 \pm 0.0150$ |
| SVC | $0.8525 \pm 0.0217$ | $0.8480 \pm 0.0223$ |
| **CNN** | **$0.9572 \pm 0.0134$** | **$0.9558 \pm 0.0148$** |
| LSTM | $0.6675 \pm 0.0262$ | $0.6546 \pm 0.0277$ |
| Transformer | $0.7739 \pm 0.0745$ | $0.7944 \pm 0.0589$ |

**TABLE II.** Classification performance of models at subject-level separation from Kumar et al. [21].

| Model | Accuracy | F1-score |
| --- | --- | --- |
| LightGBM | $0.6621 \pm 0.0352$ | $0.6566 \pm 0.0396$ |
| **Random Forest** | **$0.6823 \pm 0.0217$** | **$0.6702 \pm 0.0451$** |
| XGBoost | $0.6636 \pm 0.0274$ | $0.6533 \pm 0.0395$ |
| SVC | $0.6508 \pm 0.0572$ | $0.6276 \pm 0.1068$ |
| CNN | $0.6635 \pm 0.0529$ | $0.6415 \pm 0.0806$ |
| LSTM | $0.6170 \pm 0.0320$ | $0.6059 \pm 0.0455$ |
| Transformer | $0.6335 \pm 0.0278$ | $0.6367 \pm 0.0431$ |

The present study investigates QSVM and HQNN for detecting acute sleep deprivation using the EEG dataset of Xiang et al. Model performance is evaluated under both epoch-level and subject-level splits and compared with a classical SVM baseline.

The main contributions of this work are:

- To the best of our knowledge, this is the first application of Quantum Machine Learning Models for the detection of acute sleep deprivation from resting-state EEG signals.
- Systematic evaluation of qubit count, sample size, and scaling strategies.
- Robust validation using subject-level and epoch-level splits [21].

This work bridges QML and EEG-based neurological assessment, supporting the potential of QSVM as a complementary tool for objective sleep deprivation detection [21], [22], [28].

## II. Methodology

### A. Dataset

For this study, we used the data set of 71 participants (37 males and 34 females) between ages 17 and 23 [29]. The participants were healthy and didn’t show signs of anxiety, depression, insomnia, sudden awakening, or difficulty breathing. The participants went through two sessions, one normal sleep (NS) and one sleep deprivation (SD), with sessions counterbalanced to mitigate sequence effects. In the NS condition, participants maintained their regular sleep patterns, which were verified using sleep diaries or actigraphy. In contrast, during the SD condition, participants were required to remain awake overnight under continuous supervision, with restrictions on caffeine, alcohol, and strenuous activities. An interval of 7 days to one month was maintained between NS and SD sessions.

**Fig. 1.**
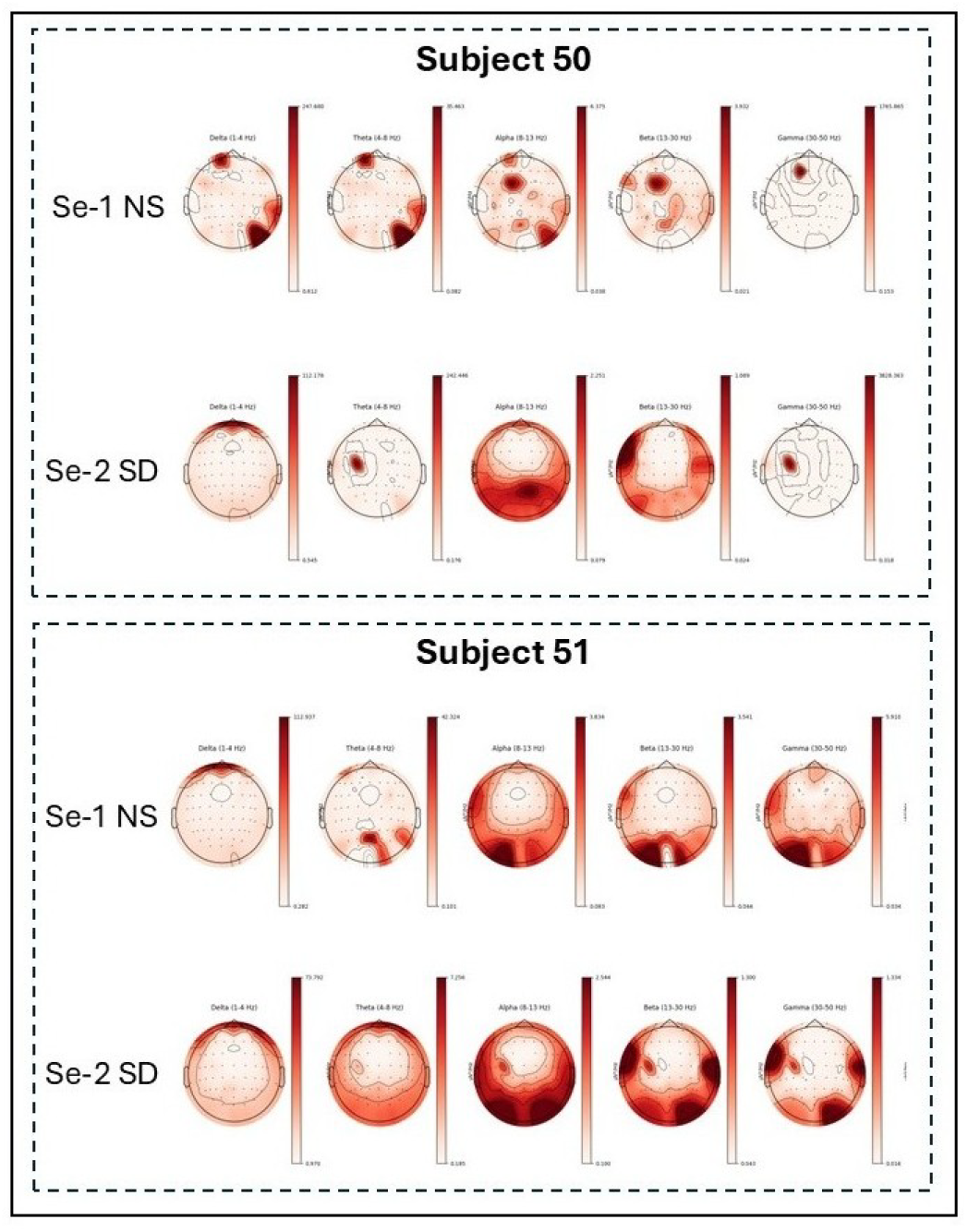
Brain heatmaps from participants 50 and 51 showing the areas of Delta, Theta, Alpha, Beta and Gamma waves for both Normal Sleep (NS) and Sleep Deprived (SD) sessions

Each session followed the same procedure. Participants first completed the Psychomotor Vigilance Test (PVT), followed by a series of state questionnaires that assess mood, anxiety, and sleepiness. The EEG data was then recorded using a 61-electrode system arranged according to the international 10–20 system, with a sampling rate of 500 Hz. Recordings included five minutes with eyes open and, for a subset of participants, an additional five minutes with eyes closed. Trait-level questionnaires were administered at the end of the session. All 71 participants completed the eyes-open condition, and 38 completed the eyes-closed condition, where each EEG recording had a duration of 5 minutes

### B. Preprocessing

All preprocessing and feature extraction procedures were implemented in Python using the MNE framework. A feature-specific electrode selection strategy was adopted to balance computational efficiency and physiological relevance. Spectral features were extracted from all available EEG channels; Hjorth parameters and theta amplitude modulation were computed from a predefined subset of frontal, central, parietal, and occipital electrodes; and coherence features were derived from a distributed set that additionally included temporal electrodes.

**Fig. 2.**
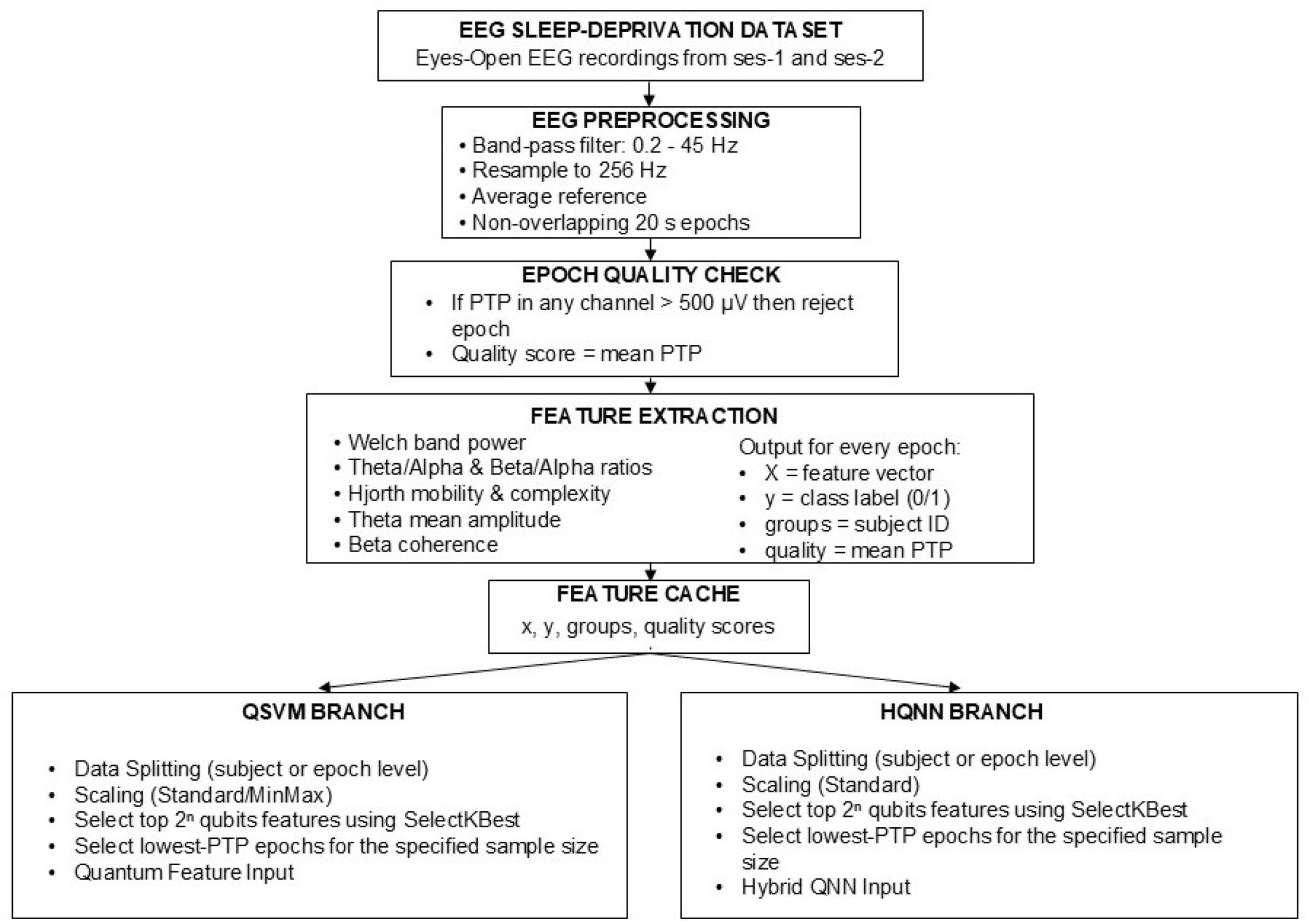
The model architecture of the proposed quantum machine learning-based sleep deprivation detection model

First, the recordings were passed through a bandpass filter from 0.2Hz to 45Hz. This was done to preserve frequency processes of neurophysiological processes. Second, to ensure a consistent rate, all recordings were resampled to 256 Hz. Third, an average reference is applied, subtracting the mean signal across all electrodes from each channel at every time point, which reduces the influence of common-mode noise and reference electrode bias.

Each cleaned signal is divided into non-overlapping 20-second windows, which balances temporal resolution with the stability of spectral estimates [30] [31]. For each window, a peak-to-peak amplitude check is performed per channel. Any epoch in which any single channel exceeds 500 microvolts peak-to-peak is discarded, a commonly used criterion for detecting non-neural artifacts such as muscle activity and electrode noise. Each surviving epoch is also assigned a quality score equal to the mean peak-to-peak amplitude across channels, which is calculated as the difference between the maximum and minimum signal amplitudes within the epoch.

This is later used to preferentially select the cleanest segments during subsampling. After preprocessing and amplitude-based artifact rejection, 1,661 usable 20-second epochs remained.

### C. Feature Extraction

Power spectral density (PSD) is calculated using Welch’s method [32] applied to each 20-second epoch across all channels simultaneously. The method segments each channel’s 20-second signal into overlapping windows of 512 samples, applies a Hanning window to each segment, computes the Fast Fourier Transform (FFT) for each windowed segment, converts the complex FFT output to power by squaring the magnitude of each frequency component, and averages the resulting periodograms across all segments to produce the final PSD estimate for that channel. This process yields a frequency resolution of 0.5 Hz and reduces variance in the spectral estimate compared to a single FFT of the entire epoch.

The extracted power spectral density is further analyzed across four standard EEG frequency bands: delta (0.5–4 Hz), theta (4–8 Hz), alpha (8–12 Hz), and beta (12–30 Hz). For each band, frequency bins within the specified range are selected using a boolean mask. The corresponding power values are then summed over the frequency axis for each channel, yielding the absolute band power per electrode.

In addition to absolute power, relative band power is also computed for each electrode. This is obtained by normalizing the band-specific power by the total power across all frequencies for that channel. Relative power provides a scale-independent measure, allowing for better comparison across subjects with varying signal amplitudes. Following band power extraction, the theta-to-alpha and the beta-to-alpha ratios are computed for the electrode. This is because theta activity increases due to sleep deprivation [33] and higher beta-band power during sleep is associated with disturbed sleep [34]

Hjorth parameters are computed for a predefined subset of electrodes (Fp1, Fp2, Fpz, Fz, Cz, Pz, and Oz). These parameters are derived in the time domain using successive finite differences of the signal, where the first-order difference approximates the first derivative and the second-order difference approximates the second derivative. The variances of the original signal, its first derivative, and its second derivative are calculated. Hjorth mobility is defined as the square root of the ratio between the variance of the first derivative and that of the original signal, providing an estimate of the signal’s mean frequency. Hjorth complexity is computed as the ratio of the mobility of the first derivative to that of the original signal, thereby reflecting the degree of frequency variation over time [35]. Additionally, to track the amplitude dynamics of frontal theta waves, an analytical signal feature was introduced. The raw signals from channels Fp1, Fp2, and Fpz were bandpass filtered to the theta frequency range (4.0–8.0 Hz). The analytical signal was then computed via the Hilbert transform, and the mean of its absolute envelope was extracted to represent the average amplitude modulation (mean AM) of frontal theta activity for each epoch.

Functional connectivity was quantified by computing coherence [36] across a set of spatially distributed electrodes covering frontal, central, parietal, occipital, and temporal regions (Fz, Cz, Pz, Oz, T7, and T8). All unique pairwise combinations of the valid electrodes are enumerated, and for each pair, coherence is computed using the same two-second sub-window length as the Welch PSD computation. The beta band frequency bins are selected using a boolean mask, and the mean coherence within the beta band is computed for that electrode pair. All pairwise beta coherence values are then averaged into a single summary feature. If fewer than two valid electrodes are found, this feature defaults to zero.

Electrode selection for different feature groups was guided by the functional characteristics of each feature type. Hjorth parameters were computed only on a subset of electrodes (Fp1, Fp2, Fpz, Fz, Cz, Pz, Oz) because they capture local temporal dynamics such as signal variance and frequency behavior, which are most informative in frontal and parietal regions known to be sensitive to sleep deprivation effects [35], [37]. In contrast, band power features were extracted across broader regions, since spectral power reflects distributed oscillatory activity and global changes in the brain state, including well-documented shifts in the theta and alpha bands during sleep deprivation [38]. Functional connectivity features were computed using a distributed and bilateral electrode set (Fz, Cz, Pz, Oz, T7, T8) to capture large-scale network synchronization, as coherence-based measures are designed to quantify inter-regional interactions and are known to reflect disrupted connectivity under sleep loss [36], [39]. This feature-specific spatial strategy ensures physiologically meaningful representation of local dynamics, global spectral activity, and network-level interactions.

For QML model training, normal sleep epochs were labeled 0 and sleep-deprived epochs were labeled 1, making this a binary classification problem. The resulting feature matrix was then passed into the quantum-model pipeline, where dimensionality reduction, normalization, and epoch subsampling were applied as described in the following subsections.

All experiments were conducted with a fixed random seed of 11. The complete dataset contained 1661 artifact-free 20-second epochs. However, because the quantum kernel requires a circuit evaluation for every pair of data points, the computational cost grows quadratically with the number of samples, making it infeasible to use the entire epoch set on classical simulators. Therefore, for both QSVM and HQNN experiments, only the cleanest epochs were retained, with subset sizes of 100, 200, and 400 epochs systematically evaluated. Within each subset, approximately two-thirds were used for training and one-third for testing, with class balancing applied independently to each split.

### D. Quantum Algorithms

#### 1) Quantum Support Vector Machine

Quantum SVM uses the quantum state space as a feature space for analyzing data. In other words, QSVM is an evolved SVM that uses quantum computing features [23]. Instead of using a normal (classical) function to measure how similar two data points are, we use a quantum version. In this quantum approach, each data point is turned into a quantum state on a set of qubits, and similarity is measured by how much these two quantum states overlap with each other.

Prior to quantum kernel computation, the high-dimensional EEG feature vectors must undergo dimensionality reduction and strict geometric preparation to satisfy the structural constraints imposed by amplitude embedding. Since a register of *n* qubits spans a Hilbert space of dimension 2^*n*^, the input classical vector must contain exactly 2^*n*^ elements, that is, 16 for the 4 qubit configuration.

To prepare the features for the quantum circuit, we first select the most useful ones and convert them into the required format. We use the ANOVA F-statistic to score each extracted feature by how well it separates the sleep-deprived class from the normal-sleep class, and then retain only the top 16 highest-scoring features. The number 16 is determined by the quantum circuit: a circuit with 4 qubits can represent 2^*n*^ = 2^4^ = 16. Finally, each 16-dimensional feature vector is divided by its Euclidean norm so that the sum of the squared amplitudes equals 1. This normalization is required for amplitude embedding, where the feature values are encoded as quantum-state probability amplitudes, whose squared magnitudes must sum to unity [40].

**Fig. 3.**
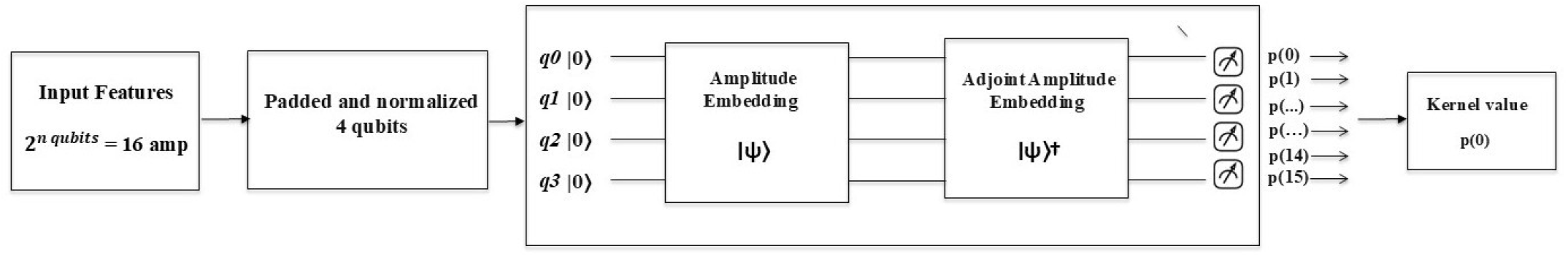
Diagram showing the architecture of of Quantum Support Vector Machine

Because computing a quantum kernel is computationally expensive, it requires running a quantum circuit for every pair of data points; the pipeline does not use the entire training and test sets. Instead, a quality-aware subsampling strategy selects the cleanest epochs. Within each class, epochs are ranked by their mean peak-to-peak amplitude, and those with the lowest values are selected first. The total subset is split approximately two-thirds for training, and one-third for testing, with the class-balanced selection applied independently to each split to prevent class imbalance from skewing the kernel computation.

The quantum kernel circuit operates on a register of *n* qubits initialized to the ground state. For any two input data points, the circuit proceeds in two sequential stages. First, amplitude embedding encodes the first normalized feature vector into the probability amplitudes of the quantum state, mapping the classical data point into a specific location in the 2^*n*^ dimensional Hilbert space. Second, the adjoint of the amplitude embedding of the second data point is applied, effectively rotating the first encoded state back through the second encoding. The probability of subsequently measuring the all-zeros computational basis state yields the quantum kernel value, which is precisely the squared modulus of the inner product between the two encoded quantum states, *k*(*x*_1_, *x*_2_) = |⟨*ϕ*(*x*_1_) |*ϕ*(*x*_2_) ⟩|^2^, ranging between zero (orthogonal) and one (identical).

The kernel is evaluated exhaustively across all required data point pairs. For training, a symmetric matrix is constructed in which each entry represents the quantum kernel value between training samples *i* and *j*. For inference, a rectangular matrix is computed between all test and training samples. These matrices fully define the similarity geometry supplied to the SVM; no raw feature vectors are passed directly to the classifier A standard SVM with a precomputed kernel is trained on the training kernel matrix and corresponding binary labels. The SVM identifies the maximum-margin hyperplane in the Hilbert space implicitly defined by the quantum kernel through quadratic programming [23]. At inference, the test kernel matrix is passed to the trained classifier, which produces binary predictions of normal sleep or sleep deprivation for each epoch, and classification accuracy is computed against the ground truth labels.

#### 2) Hybrid Quantum Neural Network

Using the same feature selection and quality-aware subsampling strategy as the QSVM pipeline, the input feature vectors were first standardized using StandardScaler and subsequently reduced to 2^*n*^ dimensions using ANOVA F-statistic-based feature selection. A classical fully-connected linear layer then projects these 2^*n*^ features onto an *n*-dimensional latent space, followed by a hyperbolic tangent activation. This bounded representation serves as the input to the quantum circuit. The quantum circuit encodes each of the *n* latent values as a rotation angle on a dedicated qubit through angle embedding. A sequence of strongly entangling layers comprising trainable single-qubit rotations interleaved with entangling operations across all qubits is subsequently applied. The number of these layers is treated as a tunable hyperparameter. After the entangling layers, the expectation value of the Pauli-Z operator is measured on each qubit, producing an *n*-dimensional vector of quantum observables [41]. These observables are passed through a final classical linear layer that reduces them to a single scalar, which is then transformed by a sigmoid activation to yield the probability of the sleep-deprived class. The entire hybrid model is trained end-to-end using binary cross-entropy loss and the Adam optimizer [42]. Early stopping with a patience of three epochs (monitored on training loss) is employed to avoid overfitting on the subsampled data. At inference, a decision threshold of 0.5 is applied to the sigmoid output to obtain binary predictions, and accuracy is computed on the held-out test subset. To ensure direct comparability with the QSVM experiments, the HQNN is evaluated across the identical experimental grid of qubit counts (2, 3, 4, and 5), subset sizes (100, 200, and 400 epochs), and validation strategies (epoch-level and subject-level splits). Additional hyperparameters specific to the HQNN learning rates (0.1, 0.01, 0.001) and number of strongly entangling layers (1, 2, 3) are systematically varied. This configuration allows quantification of how performance scales with quantum resources (qubits and circuit depth) as well as classical optimization settings. By replacing the static quantum kernel with a differentiable, parameterized circuit, the hybrid HQNN functions as an adaptive quantum feature extractor that can discover complex, data-driven transformations in the Hilbert space, potentially capturing EEG patterns that remain inaccessible to the non-trainable QSVM kernel.

**Fig. 4.**
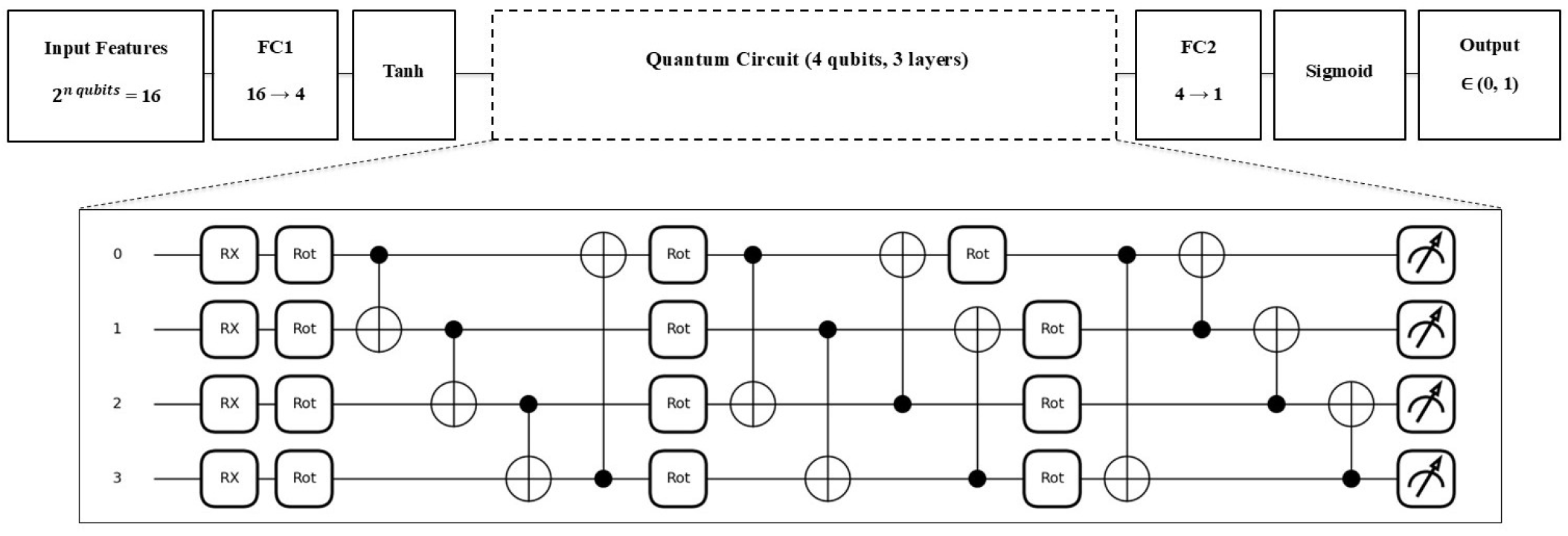
Diagram showing the architecture of Hybrid Quantum Neural Network

## III. Results

### 1) Quantum Support Vector Machine

The QSVM classifier shown in tables III and IV demonstrated promising classification performance across varying qubit counts (2–5) and sample sizes of 100, 200, and 400 epochs. The highest accuracy score for epoch-level classification of 93.75% was achieved through a qubit count of 4, 100 epochs, and minmax normalization, and for subject-level separation, a score of 75.00% was achieved through a qubit count of 5, 100 epochs, and standard normalization.

**TABLE III.** Best QSVM configurations for epoch level split.

| Qubits | Epochs | Scaler | Acc | Prec | Rec | F1 |
| --- | --- | --- | --- | --- | --- | --- |
| 2 | 100 | minmax | 0.7500 | 0.7000 | 0.8750 | 0.7778 |
| 2 | 200 | minmax | 0.6667 | 0.6774 | 0.6364 | 0.6562 |
| 2 | 400 | standard | 0.6439 | 0.8519 | 0.3485 | 0.4946 |
| 3 | 100 | minmax | 0.8125 | 0.7500 | 0.9375 | 0.8333 |
| 3 | 200 | minmax | 0.7727 | 0.8000 | 0.7273 | 0.7619 |
| 3 | 400 | standard | 0.6894 | 0.9032 | 0.4242 | 0.5773 |
| **4** | **100** | **minmax** | **0.9375** | **0.9375** | **0.9375** | **0.9375** |
| 4 | 200 | minmax | 0.8485 | 0.8710 | 0.8182 | 0.8438 |
| 4 | 400 | minmax | 0.7879 | 0.7714 | 0.8182 | 0.7941 |
| 5 | 100 | minmax | 0.8750 | 0.9286 | 0.8125 | 0.8667 |
| 5 | 200 | minmax | 0.8485 | 0.8710 | 0.8182 | 0.8438 |
| 5 | 400 | minmax | 0.7955 | 0.8197 | 0.7576 | 0.7874 |

**TABLE IV.** Best QSVM configurations for subject level split.

| Qubits | Epochs | Scaler | Acc | Prec | Rec | F1 |
| --- | --- | --- | --- | --- | --- | --- |
| 2 | 100 | standard | 0.6562 | 0.6923 | 0.5625 | 0.6207 |
| 2 | 200 | minmax | 0.6515 | 0.8125 | 0.3939 | 0.5306 |
| 2 | 400 | minmax | 0.6970 | 0.7500 | 0.5909 | 0.6610 |
| 3 | 100 | standard | 0.7188 | 0.8182 | 0.5625 | 0.6667 |
| 3 | 200 | standard | 0.7121 | 0.7692 | 0.6061 | 0.6780 |
| 3 | 400 | standard | 0.7424 | 0.7581 | 0.7121 | 0.7344 |
| 4 | 100 | standard | 0.6875 | 0.8000 | 0.5000 | 0.6154 |
| 4 | 200 | minmax | 0.7424 | 0.8333 | 0.6061 | 0.7018 |
| 4 | 400 | minmax | 0.7121 | 0.6842 | 0.7879 | 0.7324 |
| **5** | **100** | **standard** | **0.7500** | **0.8333** | **0.6250** | **0.7143** |
| 5 | 200 | standard | 0.7121 | 0.8182 | 0.5455 | 0.6545 |
| 5 | 400 | standard | 0.6970 | 0.7500 | 0.5909 | 0.6610 |

### 2) Hybrid Quantum Neural Network

The Hybrid HQNN classifier shown in tables V and VI demonstrated best classification performance across varying qubit counts, learning rates, quantum layers, and sample sizes of 100, 200, and 400 epochs. The highest accuracy score for epoch-level classification of 96.88% was achieved through a qubit count of 4, 100 epochs, learning rate of 0.1 and 3 quantum layers and for subject-level separation a score of 81.25% was achieved through a qubit count of 3, 100 epochs, learning rate of 0.01 and 3 quantum layers.

**TABLE 5.** Best HQNN configurations for epoch-level split. Here, Qrepresents the number of qubits. LR represents Learning Rate and L represents the number of layers

| Qubits | Epochs | LR | L | Acc | Prec | Rec | F1 |
| --- | --- | --- | --- | --- | --- | --- | --- |
| 2 | 100 | 0.1 | 1 | 0.8750 | 0.8750 | 0.8750 | 0.8750 |
| 2 | 200 | 0.1 | 1 | 0.7727 | 0.9091 | 0.6061 | 0.7273 |
| 2 | 400 | 0.1 | 2 | 0.6970 | 0.7955 | 0.5303 | 0.6364 |
| 3 | 100 | 0.1 | 1 | 0.9375 | 0.9375 | 0.9375 | 0.9375 |
| 3 | 200 | 0.1 | 3 | 0.8939 | 0.9643 | 0.8182 | 0.8852 |
| 3 | 400 | 0.1 | 3 | 0.7500 | 0.8000 | 0.6667 | 0.7273 |
| **4** | **100** | **0.1** | **3** | **0.9688** | **0.9412** | **1.0000** | **0.9697** |
| 4 | 200 | 0.1 | 1 | 0.8939 | 0.9062 | 0.8788 | 0.8923 |
| 4 | 400 | 0.1 | 3 | 0.8182 | 0.7763 | 0.8939 | 0.8310 |
| 5 | 100 | 0.1 | 3 | 0.9375 | 0.9375 | 0.9375 | 0.9375 |
| 5 | 200 | 0.1 | 1 | 0.9091 | 0.8857 | 0.9394 | 0.9118 |
| 5 | 400 | 0.01 | 2 | 0.7955 | 0.9756 | 0.6061 | 0.7477 |

**TABLE 6.** Best HQNN Configurations for subject-level split. Here, Qrepresents the number of qubits. LR represents learning Rate and L represents the number of layers

| Qubits | Epochs | LR | L | Acc | Prec | Rec | F1 |
| --- | --- | --- | --- | --- | --- | --- | --- |
| 2 | 100 | 0.1 | 3 | 0.6250 | 0.7000 | 0.4375 | 0.5385 |
| 2 | 200 | 0.1 | 2 | 0.7121 | 0.8182 | 0.5455 | 0.6545 |
| 2 | 400 | 0.1 | 1 | 0.7652 | 0.8431 | 0.6515 | 0.7350 |
| **3** | **100** | **0.01** | **3** | **0.8125** | **1.0000** | **0.6250** | **0.7692** |
| 3 | 200 | 0.01 | 1 | 0.7727 | 0.9500 | 0.5758 | 0.7170 |
| 3 | 400 | 0.01 | 1 | 0.7652 | 0.9070 | 0.5909 | 0.7156 |
| 4 | 100 | 0.1 | 3 | 0.7188 | 1.0000 | 0.4375 | 0.6087 |
| 4 | 200 | 0.01 | 3 | 0.7424 | 0.8333 | 0.6061 | 0.7018 |
| 4 | 400 | 0.1 | 3 | 0.7045 | 0.6901 | 0.7424 | 0.7153 |
| 5 | 100 | 0.01 | 1 | 0.7812 | 0.9091 | 0.6250 | 0.7407 |
| 5 | 200 | 0.1 | 1 | 0.7576 | 0.8400 | 0.6364 | 0.7241 |
| 5 | 400 | 0.1 | 2 | 0.7045 | 0.6800 | 0.7727 | 0.7234 |

## IV. Conclusion and future work

This study explores the use of a Quantum Support Vector Machine (QSVM) and Hybrid Quantum Neural Networks (HQNN) for the detection of acute sleep deprivation using resting state EEG signals. The results demonstrate that HQNN achieves competitive performance, with accuracy reaching 81.25% under subject-level evaluation and 96.88% under epoch-level evaluation, greatly improving upon previously reported results. Although higher accuracy was observed under epoch-level evaluation, this setting is less reliable due to potential data overlap. The subject-level results, therefore, provide a more realistic assessment of model performance and highlight the ability of QSVM and Hybrid HQNN to generalize across individuals.

**Fig. 5.**
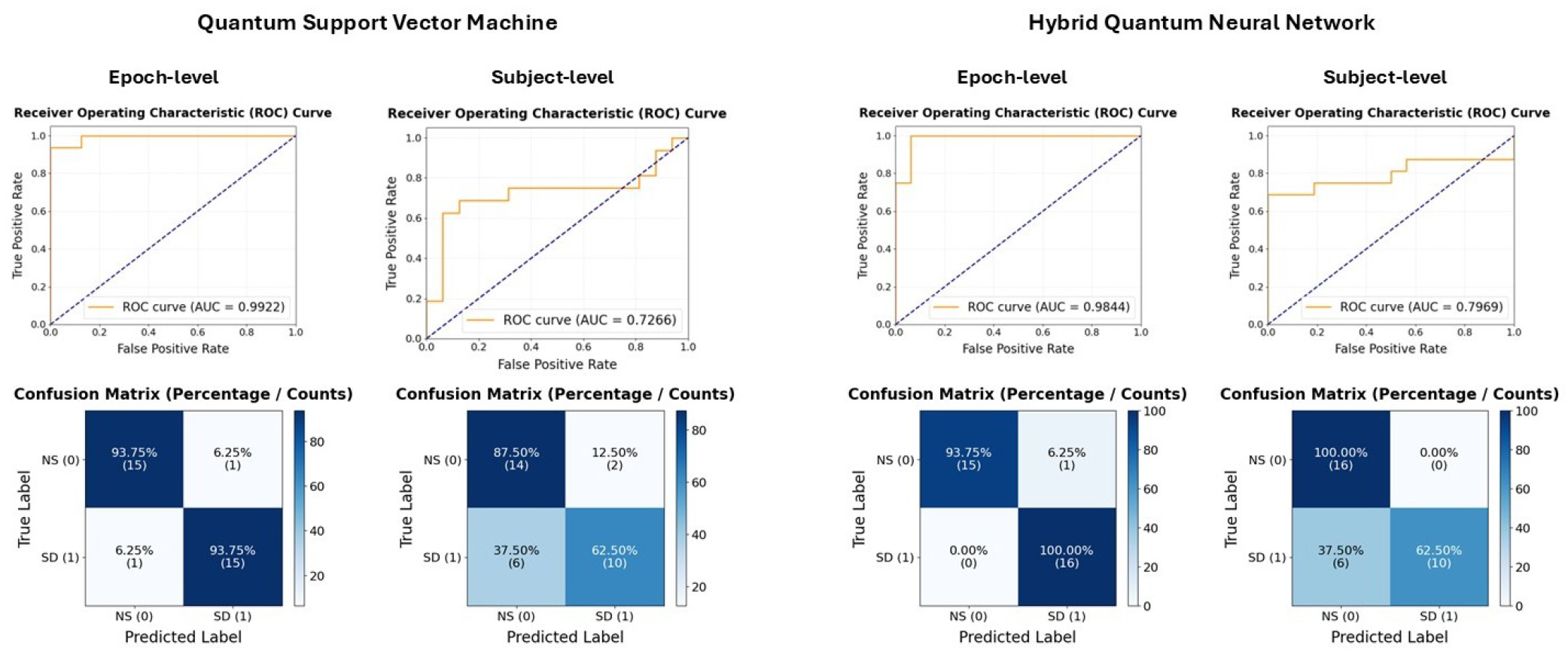
Confusion matrix and ROC graphs of Quantum Support Vector Machine and Hybrid Quantum Neural Network for both subject-level and epoch-level separation

Despite these promising findings, the study is limited by the relatively small dataset and the computational constraints associated with quantum kernel evaluation. Furthermore, current results are obtained using simulated quantum environments, and performance may vary on real quantum hardware. Future work should focus on scaling the approach to larger and more diverse datasets, exploring alternative quantum feature maps, and evaluating hybrid quantum-classical models. Investigating additional EEG features and improving cross-subject generalization will also be important for practical applications. Ultimately, this work supports the growing potential of quantum machine learning in biomedical signal analysis and suggests that QSVM and Hybrid HQNN can serve as a promising tool for the detection of cognitive disorders.

## V. Use of Generative AI

Generative AI tools, including ChatGPT (OpenAI, GPT-5.3), were used solely to improve the clarity, grammar, and readability of this manuscript. All content was reviewed and validated by the authors to ensure accuracy and compliance with academic standards.

## References

[1] V. V. Vyazovskiy, U. Olcese, E. C. Hanlon, Y. Nir, C. Cirelli, and G. Tononi, “Local sleep in awake rats,” Nature, vol. 472, pp. 443–447, 2011.

[2] C.-S. Hung, S. Sarasso, F. Ferrarelli, B. Riedner, M. F. Ghilardi, C. Cirelli, and G. Tononi, “Local experience-dependent changes in the wake EEG after prolonged wakefulness,” Sleep, vol. 36, pp. 59–72, 2013.

[3] K. C. Kayser, V. A. Puig, and J. R. Estepp, “Predicting and mitigating fatigue effects due to sleep deprivation: a review,” Frontiers in Neuroscience, vol. 16, p. 930280, 2022.

[4] G. Bernardi, F. Siclari, X. Yu, C. Zennig, M. Bellesi, E. Ricciardi et al., “Neural and behavioral correlates of extended training during sleep deprivation in humans: evidence for local, task-specific effects,” Journal of Neuroscience, vol. 35, pp. 4487–4500, 2015.

[5] R. M. Brossoit, T. L. Crain, J. J. Leslie, L. B. Hammer, D. M. Truxillo, and T. E. Bodner, “The effects of sleep on workplace cognitive failure and safety,” Journal of Occupational Health Psychology, vol. 24, pp. 411–422, 2019.

[6] W. D. S. Killgore, “Effects of sleep deprivation on cognition,” Progress in Brain Research, vol. 185, pp. 105–129, 2010.

[7] M. M. Mitler and J. C. Miller, “Methods of testing for sleepiness,” Behavioral Medicine, vol. 21, pp. 171–183, 1996.

[8] J. Lian, L. Xu, T. Song, Z. Peng, Z. Zhang, X. An et al., “Reduced resting-state EEG power spectra and functional connectivity after 24 and 36 hours of sleep deprivation,” Brain Sciences, vol. 13, p. 949, 2023.

[9] Z. Liu, Y. Zhou, C. Hao, and N. Ma, “Alteration in neural oscillatory activity and phase-amplitude coupling after sleep deprivation: evidence for impairment and compensation effects,” Journal of Sleep Research, vol. 34, p. e14264, 2025.

[10] C. Cajochen, V. Knoblauch, K. Kräuchi, C. Renz, and A. Wirz-Justice, “Dynamics of frontal EEG activity, sleepiness and body temperature under high and low sleep pressure,” NeuroReport, vol. 12, pp. 2277–2281, 2001.

[11] I. M. Verweij, N. Romeijn, D. J. Smit, G. Piantoni, E. J. Van Someren, and Y. D. van der Werf, “Sleep deprivation leads to a loss of functional connectivity in frontal brain regions,” BMC Neuroscience, vol. 15, p. 88, 2014.

[12] S. Zhao, F. Long, X. Wei, X. Ni, H. Wang, and B. Wei, “Evaluation of a single-channel EEG-based sleep staging algorithm,” International Journal of Environmental Research and Public Health, vol. 19, p. 2845, 2022.

[13] M. M. Monowar, S. N. Nobel, M. Afroj, M. A. Hamid, M. Z. Uddin, M. M. Kabir et al., “Advanced sleep disorder detection using multi-layered ensemble learning and advanced data balancing techniques,” Frontiers in Artificial Intelligence, vol. 7, p. 1506770, 2025.

[14] Y. Wang, S. Ye, Z. Xu, Y. Chu, J. Zhang, and W. Yu, “Research on sleep staging based on support vector machine and extreme gradient boosting algorithm,” Nature and Science of Sleep, vol. 16, pp. 1827–1847, 2024.

[15] L. A. Wang, R. Kern, E. Yu, S. Choi, and J. Q. Pan, “IntelliSleepScorer, a software package with a graphic user interface for automated sleep stage scoring in mice based on a light gradient boosting machine algorithm,” Scientific Reports, vol. 13, p. 4275, 2023.

[16] R. Jain and R. A. Ganesan, “Effective diagnosis of various sleep disorders by LEE classifier: LightGBM-EOG-EEG,” IEEE Journal of Biomedical and Health Informatics, vol. 29, pp. 2581–2588, 2025.

[17] U. Kumari, P. Kora, K. Meenakshi, K. Swaraja, T. Padma, A. K. Panigrahy et al., “Feature extraction and detection of obstructive sleep apnea from raw EEG signal,” in Proceedings of the International Conference on Innovative Computing and Communications (ICICC 2019), vol. 1. Springer Singapore, 2020, pp. 425–433.

[18] Z. Zhang, Y. Xue, A. Slowik, and Z. Yuan, “SLE-CNN: a novel convolutional neural network for sleep stage classification,” Neural Computing and Applications, vol. 35, pp. 17 201–17216, 2023.

[19] L. Zhuang, M. Dai, Y. Zhou, and L. Sun, “Intelligent automatic sleep staging model based on CNN and LSTM,” Frontiers in Public Health, vol. 10, p. 946833, 2022.

[20] C. Wan, M. C. Nnamdi, W. Shi, B. Smith, C. Purnell, and M. D. Wang, “Advancing sleep disorder diagnostics: a transformer-based EEG model for sleep stage classification and OSA prediction,” IEEE Journal of Biomedical and Health Informatics, vol. 29, pp. 878–886, 2025.

[21] D. Kumar, A. Narayan, and S. Lalgudi Ganesan, “An EEG-based machine learning framework for diagnosing acute sleep deprivation,” Frontiers in Physiology, vol. 16, p. 1668129, 2025.

[22] Y. Gujju, A. Matsuo, and R. Raymond, “Quantum machine learning on near-term quantum devices: Current state of supervised and unsupervised techniques for real-world applications,” 2023, arXiv:2307.00908.

[23] V. Havlíček, A. D. Córcoles, K. Temme, A. W. Harrow, A. Kandala, J. M. Chow, and J. M. Gambetta, “Supervised learning with quantum-enhanced feature spaces,” Nature, vol. 567, no. 7747, pp. 209–212, 2019.

[24] Y. Kumar et al., “Heart failure detection using quantum-enhanced machine learning and traditional machine learning techniques for Internet of Artificially Intelligent Medical Things,” Wireless Communications and Mobile Computing, vol. 2021, p. e1616725, 2021.

[25] H. Gupta, H. Varshney, T. K. Sharma, N. Pachauri, and O. P. Verma, “Comparative performance analysis of quantum machine learning with deep learning for diabetes prediction,” Complex & Intelligent Systems, vol. 8, pp. 3073–3087, 2022.

[26] Z. Ozpolat and M. Karabatak, “Performance evaluation of quantum-based machine learning algorithms for cardiac arrhythmia classification,” Diagnostics, vol. 13, p. 1099, 2023.

[27] A. Padha and A. K. Sahoo, “Quantum enhanced machine learning for unobtrusive stress monitoring,” in Proceedings of the 2022 Fourteenth International Conference on Contemporary Computing (IC3-2022). ACM, 2022, pp. 476–483.

[28] G. Aksoy, G. Cattan, S. Chakraborty, and M. Karabatak, “Quantum machine-based decision support system for the detection of schizophrenia from EEG records,” Journal of Medical Systems, vol. 48, no. 1, p. 29, 2024.

[29] C. Xiang, X. Fan, D. Bai, K. Lv, and X. Lei, “A resting-state eeg dataset for sleep deprivation,” Scientific Data, vol. 11, p. 427, 2024.

[30] J. Möcks and T. Gasser, “How to select epochs of the eeg at rest for quantitative analysis,” Electroencephalography and Clinical Neurophysiology, vol. 58, pp. 89–92, 1984.

[31] M. N. Mohsenvand, M. R. Izadi, and P. Maes, “Contrastive representation learning for electroencephalogram classification,” in Proceedings of the Machine Learning for Health NeurIPS Workshop, E. Alsentzer, M. B. A. McDermott, F. Falck, S. K. Sarkar, S. Roy, and S. L. Hyland, Eds., vol. 136, 2020, pp. 238–253.

[32] P. D. Welch, “The use of fast fourier transform for the estimation of power spectra: A method based on time averaging over short, modified periodograms,” IEEE Transactions on Audio and Electroacoustics, vol. 15, pp. 70–73, 1967.

[33] C. Cajochen, R. Foy, and D. J. Dijk, “Frontal predominance of a relative increase in sleep delta and theta eeg activity after sleep loss in humans,” Sleep Research Online, vol. 2, no. 3, pp. 65–69, 1999.

[34] T. B. J. Kuo, C.-Y. Chen, Y.-C. Hsu, and C. C. H. Yang, “Eeg beta power and heart rate variability describe the association between cortical and autonomic arousals across sleep,” Autonomic Neuroscience, vol. 194, pp. 32–37, 2016.

[35] B. Hjorth, “Eeg analysis based on time domain properties,” Electroencephalography and Clinical Neurophysiology, vol. 29, no. 3, pp. 306–310, 1970.

[36] P. L. Nunez, R. Srinivasan, A. F. Westdorp, R. S. Wijesinghe, D. M. Tucker, R. B. Silberstein et al., “Eeg coherency: I. statistics, reference electrode, volume conduction, laplacians, cortical imaging, and interpretation at multiple scales,” Electroencephalography and Clinical Neurophysiology, vol. 103, pp. 499–515, 1997.

[37] N. e. a. Al-Qazzaz, “Automated eeg signal analysis using hjorth parameters for brain-state classification,” Biomedical Signal Processing and Control, 2017.

[38] W. Klimesch, “Eeg alpha and theta oscillations reflect cognitive and memory performance,” Brain Research Reviews, vol. 29, pp. 169–195, 1999.

[39] F. e. a. Vecchio, “Functional connectivity alterations in sleep deprivation,” Human Brain Mapping, 2017.

[40] M. A. Nielsen and I. L. Chuang, Quantum Computation and Quantum Information, 1st ed. Cambridge: Cambridge University Press, 2000.

[41] M. Schuld, V. Bergholm, C. Gogolin, J. Izaac, and N. Killoran, “Evaluating analytic gradients on quantum hardware,” Physical Review A, vol. 99, no. 3, p. 032331, 2019.

[42] D. P. Kingma and J. Ba, “Adam: A method for stochastic optimization,” arXiv preprint arXiv:1412.6980, 2014, published as a conference paper at ICLR 2015.

